# Experimental erosion of microbial diversity decreases soil CH_4_ consumption rates

**DOI:** 10.1101/2020.03.24.003657

**Authors:** Elvira Schnyder, Paul L.E. Bodelier, Martin Hartmann, Ruth Henneberger, Pascal A. Niklaus

## Abstract

Biodiversity-ecosystem functioning (BEF) experiments have predominantly focused on communities of higher organisms, in particular plants, with comparably little known to date about the relevance of biodiversity for microbially-driven biogeochemical processes. Methanotrophic bacteria play a key role in Earth’s methane (CH_4_) cycle by removing atmospheric CH_4_ and reducing emissions from methanogenesis in wetlands and landfills. Here, we used a dilution-to-extinction approach to simulate diversity loss in a methanotrophic landfill cover soil community. Combining analyses of CH_4_ flux and community structure, we found a linear decrease of CH_4_ oxidation rates with the number of taxonomic units lost. This effect was independent of community size, consistent over the three-month study, and occurred in relatively diverse communities, challenging the notion of high functional redundancy mediating high resistance to diversity erosion in natural microbial systems. The effects we report resemble the ones for higher organisms, suggesting that BEF-relationships are universal across taxa and spatial scales.

## Introduction

Soil microbes modulate Earth’s climate by mediating fluxes of major greenhouse gases including methane (CH_4_) [1]. CH_4_ occurs at lower atmospheric volume mixing ratios than CO_2_ but exerts much larger radiative effects on a per-molecule basis so that the anthropogenic warming effect of CH_4_ is on par with that of CO_2_ on a decadal time scale [2]. Atmospheric CH_4_ loads also are of concern because they have shown an accelerated rise in recent years but the underlying drivers are not fully understood [3].

Soil microbes drive two key CH_4_ transformations: methanogenesis and CH_4_ oxidation. Methanogenic archaea produce CH_4_ under anaerobic conditions, a process that is quantitatively important in natural wetlands, rice paddies, and landfills [4]. Obligate aerobic methanotrophic bacteria oxidize CH_4_ in terrestrial soils, with less important contributions by other bacteria (e.g. ammonia oxidizing bacteria [5], and possibly also archaea under anaerobic conditions [6, 7]). CH_4_ oxidation by methanotrophs thus constitutes a biogenic sink that removes CH_4_ from the atmosphere. In wetlands and landfills, methanotrophs also remove CH_4_ produced by methanogenic archaea, and functionally thus act as a “biofilter” that eliminates a large fraction of potential soil CH_4_ emissions [8]. In these systems, changes in soil-atmosphere CH_4_ fluxes may therefore be mediated by changes in rates of CH_4_ oxidation, in rates of methanogenesis, or in both.

Over the past decades, experimental [9], observational [10], and theoretical studies [11] have demonstrated that biodiversity is critical for the provisioning of a wide range of ecosystem services. Most biodiversity experiments to date have focused on plant-related functions such as primary productivity. They generally revealed effects that were “positive decelerating”, i.e. more diverse communities performed better but the additional benefit mediated by extra species diminished when communities already contained a high number of species. These positive biodiversity-ecosystem functioning (BEF) relationships have been attributed to three groups of mechanisms. First, diverse communities are more likely to contain highly productive species, a phenomenon also known as “selection probability effect” [12, 13]. Second, BEF relationships can arise from niche differentiation that allows more diverse communities to more fully acquire essential resources; there is little doubt that such complementarity is important, but to date remarkably little progress has been made in identifying the specific mechanisms and resources concerned [14–16]. Productivity can also be promoted by complementary biotic interactions which may reduce negative impacts of pathogens and consumers if these are host-specific and their activity density-dependent [17]. Third, species can modify the environmental conditions in a way that favors the growth of another species, a mechanism called facilitation [18].

Plants are major drivers of the global carbon cycle. They have evolved a fascinating diversity of organisms that are adapted to a wide range of environments. Nevertheless, they are metabolically relatively uniform and drive essentially the same suite of biogeochemical processes. In contrast, soil microbes drive a plethora of chemically extremely diversified ecosystem functions. The ecology and life-history of microbes differ from the ones of plants and animals, potentially giving rise to divergent mechanisms underpinning microbial BEF relationships [19]. The diversity of microorganisms in soils also is extreme, exceeding the one found in higher organisms by orders of magnitude [20–23]. This supports the idea that microbial communities are less susceptible to diversity loss due to high functional redundancy. On the other hand, soils are extremely heterogeneous at the micro-scale, and some key microbial functions are carried out by phylogenetically relatively narrow groups of microbes. Aerobic CH_4_ oxidation in soils, for example, is driven by bacteria that belong to either the Gammaproteobacteria (type I methanotrophs), Alphaproteobacteria (type II methanotrophs), or Verrucomicrobia (type III methanotrophs) [24, 25]. In typical habitats, their diversity rarely exceeds a few dozen strains. It may thus well be that CH_4_ oxidation rates by these communities critically depend on the number of species present. However, experimental microbial BEF research has only recently begun to emerge [26–32] and little is understood to date on how microbe-mediated ecosystem functions depend on the diversity of the respective guilds.

Here, we analyze diversity effects on CH_4_ consumption in microbial communities extracted from the cover soil of a landfill that had accumulated several million m^3^ of municipal solid waste until the 1990s. In such cover soils, methanotrophs oxidize CH_4_ at high rates, thereby filtering most CH_4_ produced in the course of waste degradation [33, 34]. Nevertheless, landfills remain a major global anthropogenic source of CH_4_, and any temporary or permanent reduction in methanotrophic activity in cover soils can trigger massive increases in net CH_4_ emissions. To test whether methanotrophic activity in such communities is impaired by diversity loss, we created communities that spanned a gradient in the diversity of taxonomic units using a dilution-to-extinction approach. Diversity erosion by dilution allowed us to manipulate the diversity of communities with microbial members that are difficult to isolate and cultivate. It further had the advantage that the resulting diversity gradients mimicked a realistic extinction scenario where rare species were more likely to go extinct [35]. The dilutions ranged from 10^−1^ over 10^−3^, 10^−4^, 10^−5^ to 10^−7^, and were replicated in three independent extinction series. We inoculated microcosms with these dilutions, pre-incubated the microcosms to equalize community sizes, and then incubated these under headspace average CH_4_ concentrations of 7400 μmol mol^−1^ for nearly three months, at a temperature that was either constant at 15°C or which oscillated diurnally between 10 and 20°C. Each treatment combination was replicated 4 times, resulting in a total of 120 microcosms (3 series × 5 dilutions × 2 temperatures × 4 replicates). Throughout the experiment, we quantified methanotrophic activity by assessing CH_4_ consumption and CO_2_ production, with the latter also including respiration by non-methanotrophs. Using quantitative PCR (qPCR), we determined methanotrophic community sizes and growth as abundance of a key functional gene, *pmoA,* which encodes for a subunit of the particulate methane mono-oxygenase enzyme found in most methanotrophs. Total bacterial abundance was determined by ribosomal gene abundance (16S rRNA subunit). Finally, we determined the composition of the experimental communities by high-throughput amplicon sequencing of ribosomal (16S rRNA) and functional (*pmoA*) gene fragments.

Our aim was to test whether diversity loss affects methanotroph community functioning. We further were interested whether phylogenetic diversity is a good predictor of ecosystem functioning. Ultimately, BEF relationships arise from differences in functional traits among species, and phylogenetic diversity might serve as a proxy for functional differences if the relevant traits are preserved through evolutionary time [36]. Finally, we asked whether diversity effects were larger in a fluctuating environment (diurnal temperature cycle) that might provide a larger environmental niche space and allow for functional complementarity among community members.

## Results

### Community diversity

We extracted DNA from microcosms after 0, 31, 58 and 86 days (end of the experiment) and sequenced the 16S rRNA gene (V1—V2) and the *pmoA* gene using the Illumina MiSeq platform. The 11·10^6^ 16S rRNA gene and 14·10^6^ *pmoA* sequences clustered into 2697 non-methanotrophic bacterial and 58 methanotrophic OTUs (Methods). Using OTU abundances, we determined the diversity metrics richness (S), Shannon index (H), and phylogenetic diversity (PD), which were statistically highly significantly related to dilution level (Fig. 1; F_1,10_=55—274 and P=2·10^−5^—10^−8^, depending on gene and diversity metric). Importantly, the established diversity gradient was maintained throughout the entire incubation, with similarly significant effects after 86 days. The realized diversity gradient was statistically independent of the temperature treatment.

**Figure 1.**
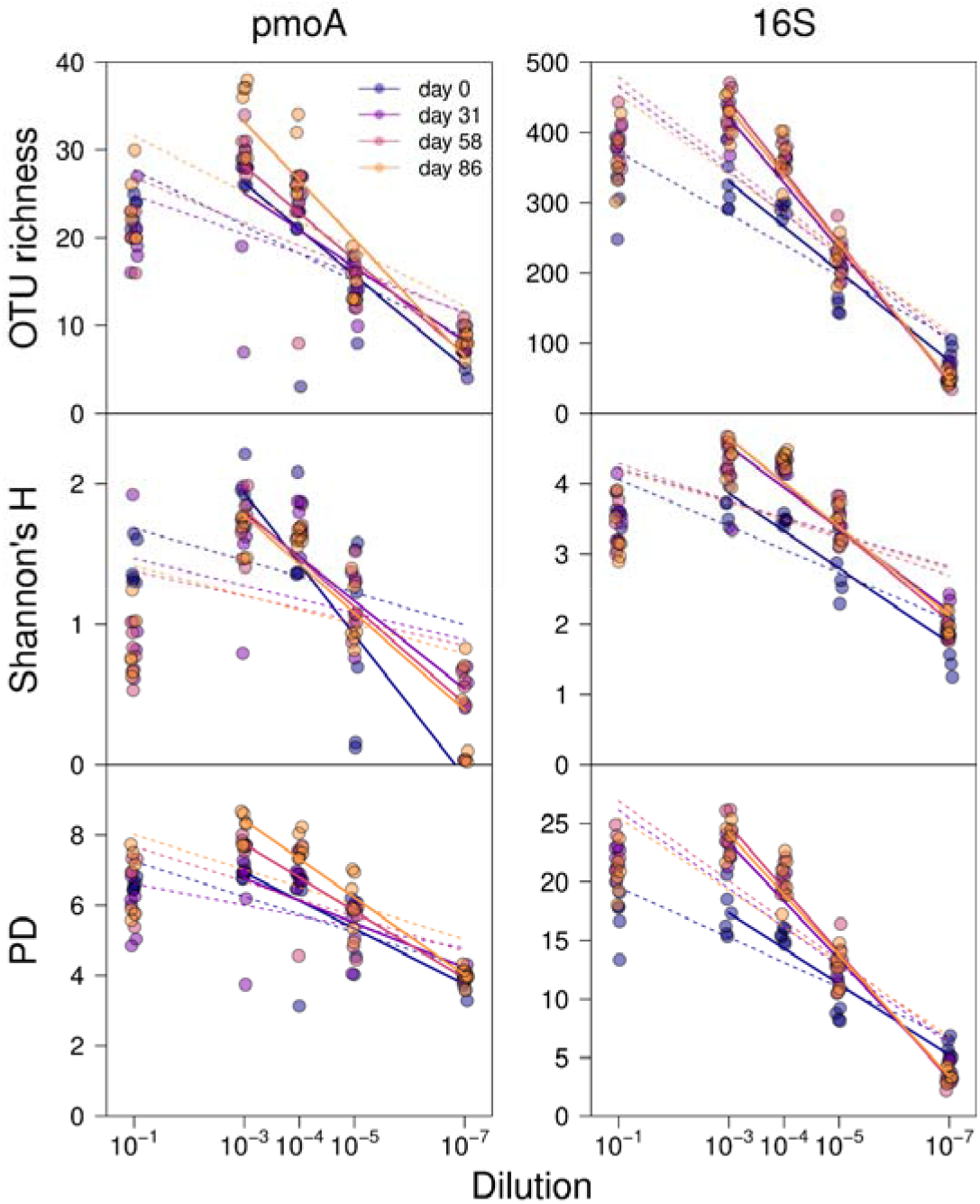
Effects of dilution on OTU richness, Shannon diversity (H), and phylogenetic diversity (PD) in dependence of dilution level and sampling date. Diversity metrics are based on rarefied sequence sets. Open symbols indicate samples for which no rarefaction was possible because sequence numbers were too low; these samples were not included in the computation of regression lines. DNA extraction was incomplete in the least diluted samples (10^−1^) where a precipitate formed. Linear regressions therefore are provided excluding (solid lines) and including (dashed liens) the 10^−1^ dilution.

### Community composition

At the end of the experiment, the most common non-methanotrophic bacterial taxa were Proteobacteria, Bacteroidetes, Actinobacteria and Verrucomicrobia. Type Ia methanotrophs, in particular *Methylobacter,* dominated the methanotroph community (Fig. 2). Across all dilutions, the proportion of type Ia methanotrophs decreased with time, with opposite effects on type IIa methanotrophs (Fig. 2). Permutational MANOVA (Methods) with Benjamini-Hochberg adjustment to a false discovery rate of 5% did not reveal any effects of dilution on non-methanotrophic bacterial community composition, at all taxonomic levels investigated (phyllum, class, order, family, genus, OTU). Conversely, low to intermediate dilution levels were characterized by high type Ia methanotrophs abundances, in particular of *Methylobacter* and *Methylosarcina.*

**Figure 2.**
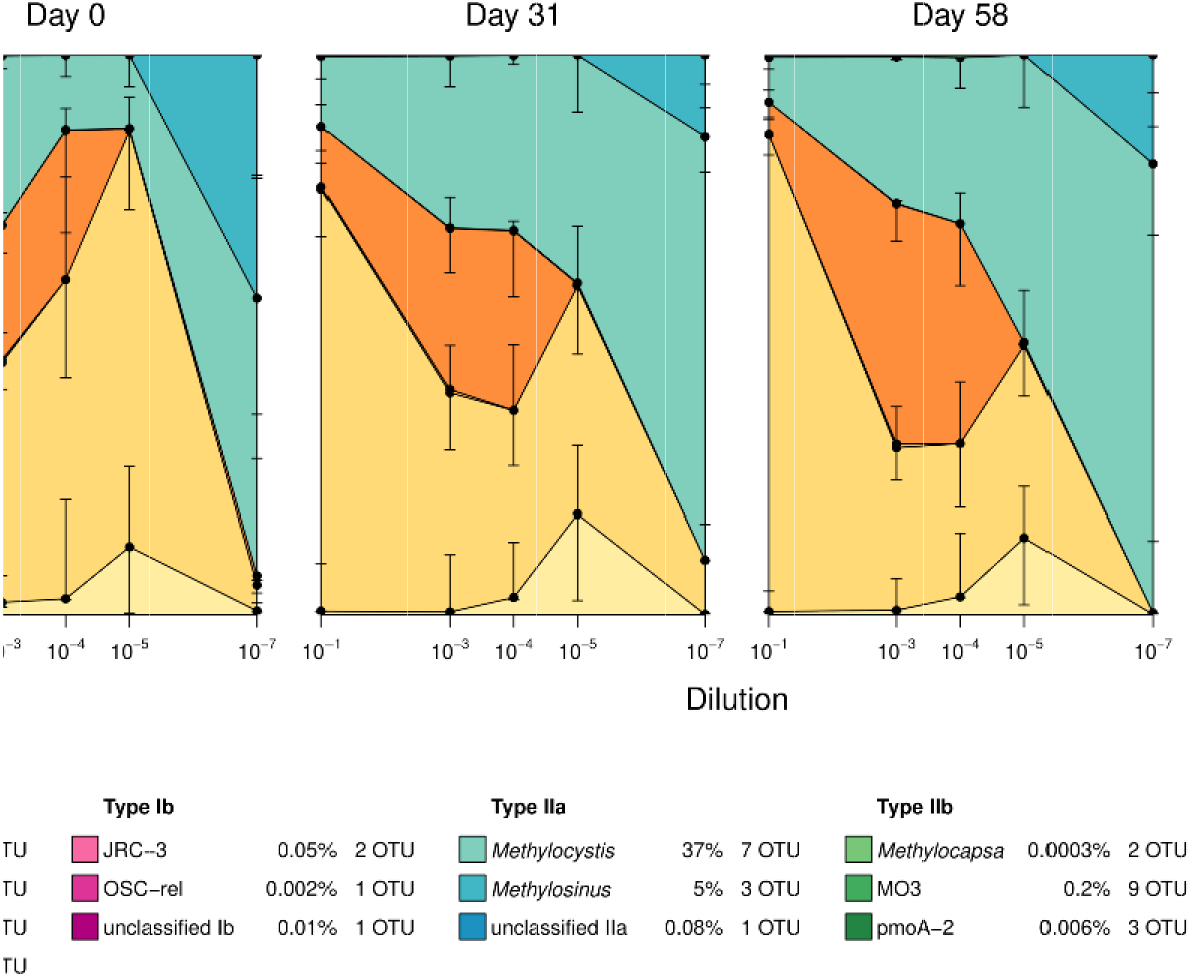
Effects of dilution on methanotroph community composition through time. The legend shows the average fractional abundance of all taxa across all sampling dates and dilution levels, together with the number of OTUs. Error bars show uncertainties for the different groups (1 s.e.) and are directed inwards.

At high dilution, the methanotroph community was almost exclusively composed of type IIa methanotrophs of the genera *Methylocystis* and *Methylosinus* (Fig. 2). The temperature treatments did not significantly affect the relative abundance of any taxa, neither for *pmoA* nor for 16S rRNA gene sequences. Principle coordinate analysis (PCoA) of Bray-Curtis dissimilarities among communities revealed that composition changed relatively little with time but that the serial dilutions resulted in a consistent trajectory in ordination space, independent of the applied temperature treatment (SI Fig. 1).

### Community size

Quantitative PCR (qPCR) analysis revealed that 16S rRNA gene copy numbers decreased in the course of the experiment but were statistically independent of dilution and temperature treatments (Fig. 3b, d). Averaged over the experiment, *pmoA* copy numbers also were statistically independent of dilution (Fig. 3a). However, at the beginning of the experiment (day 0), *pmoA* copy numbers were approximately two orders of magnitude lower in the highest dilution (10^−7^) compared to less diluted samples (Fig. 3c). This effect had disappeared at the second sampling on day 31. qPCR data for *pmoA* were very variable, with equivocal findings depending on whether data were inspected on a *log-scale* (which accounts for the exponential nature of the PCR process) or on an untransformed scale (which is a necessity to test for biodiversity effects, see Methods). The *log*-transformed data indicated a trend for communities in the high dilution treatments to grow through time, whereas the least diluted communities tended to decrease in size.

**Figure 3.**
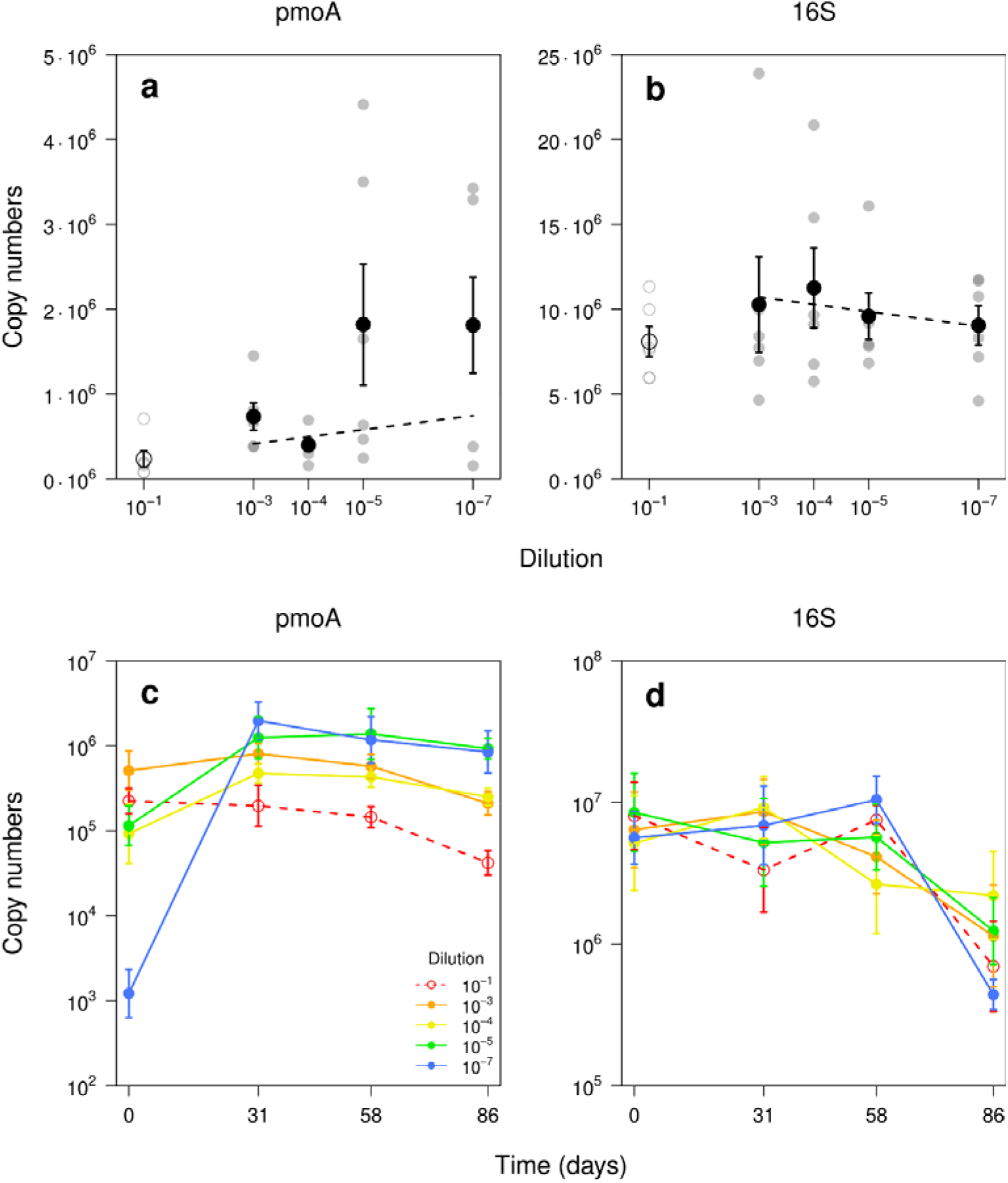
Bacterial and methanotroph community size, determined as (a,c) *pmoA* and (b,d) 16S rRNA gene copy numbers. (a,b) Gene copy numbers in dependence of applied dilution treatment. Grey symbols indicate averages across 4 sampling dates for each community composition × temperature treatment combination (3 dilution series × 5 dilution levels × 2 temperatures). Dashed lines show linear regressions accounting for heterogeneous variance among dilution levels. Note that the copy numbers are shown on a linear scale, yielding horizontal regression lines under the null hypothesis (see methods). Black symbols indicate mean copy numbers for each dilution level, with error bars indicating standard errors (n=6). (c,d) Gene copy numbers in dependence of sampling date and dilution (mean ± s.e.; n=6). Data are shown on a logarithmic scale, which accounts for the exponential error distribution introduced by the PCR process but is not suited to test for effects of diversity reductions by dilution (see methods).

### CH_4_ consumption and CO_2_ release rates

CH_4_ consumption, determined by gas chromatography of headspace samples, decreased 1.9-fold with diversity loss along the dilution gradient (P < 0.001, Fig. 4, averages over all samplings). Net CH_4_ consumption was significantly related to *pmoA*-based methanotrophic OTU richness, the Shannon diversity index H, and phylogenetic diversity PD (Fig. 4). CH_4_ consumption rates were higher initially and stabilized during the first 20 days, except for the highest dilution (10^−7^) which showed no initial activity and then overshot before also stabilizing around day 30 (Fig. 5). Towards the end of the experiment (days 60—86), diversity effects even grew larger, resulting in a 7.5-fold change from the 10^−1^ to the 10^−7^ dilution, with rates of 105±9 and 14±3 μmol CH_4_ d^−1^ microcosm^−1^, respectively. CO_2_ production very closely mirrored CH_4_ consumption (r = 0.86, Pearson’s product moment correlation, P < 0.001; data not shown). The temperature treatment did not affect net CH_4_ consumption.

**Figure 4.**
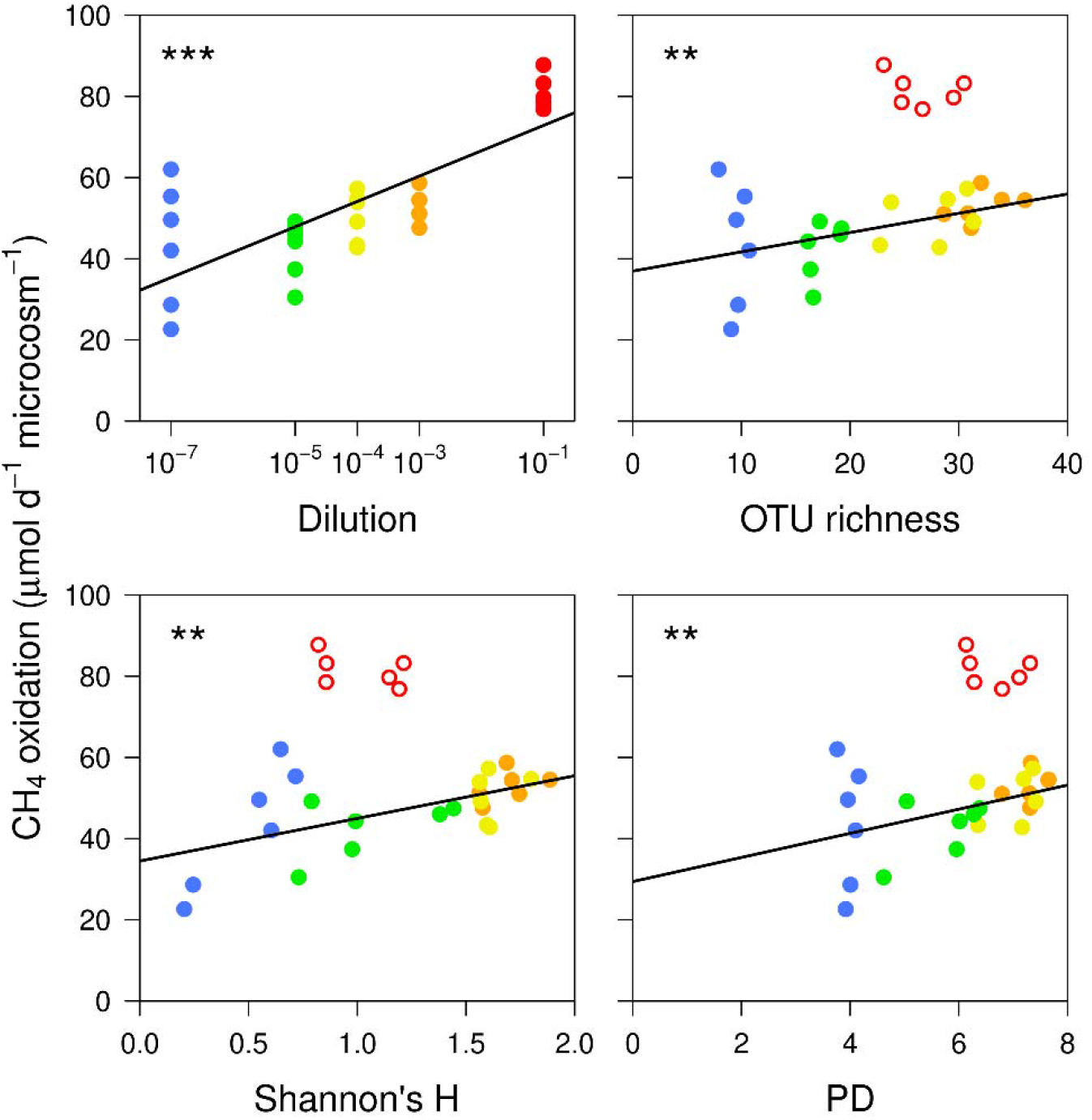
Net CH_4_ consumption rates of microcosms in dependence of dilution level and methanotrophic diversity. [*pmoA*-based OTU richness, Shannon index (H), and phylogenetic diversity (PD)]. Symbols show means for each microcosm (n=3 replicates × 2 temperature treatments per dilution level; temperature: n.s.). Samples with 10^−1^ dilution are excluded from statistical tests (indicated by open symbols) because of incomplete DNA extraction due to precipitate. *** P<0.001; ** P<0.01; * P<0.05.

**Figure 5.**
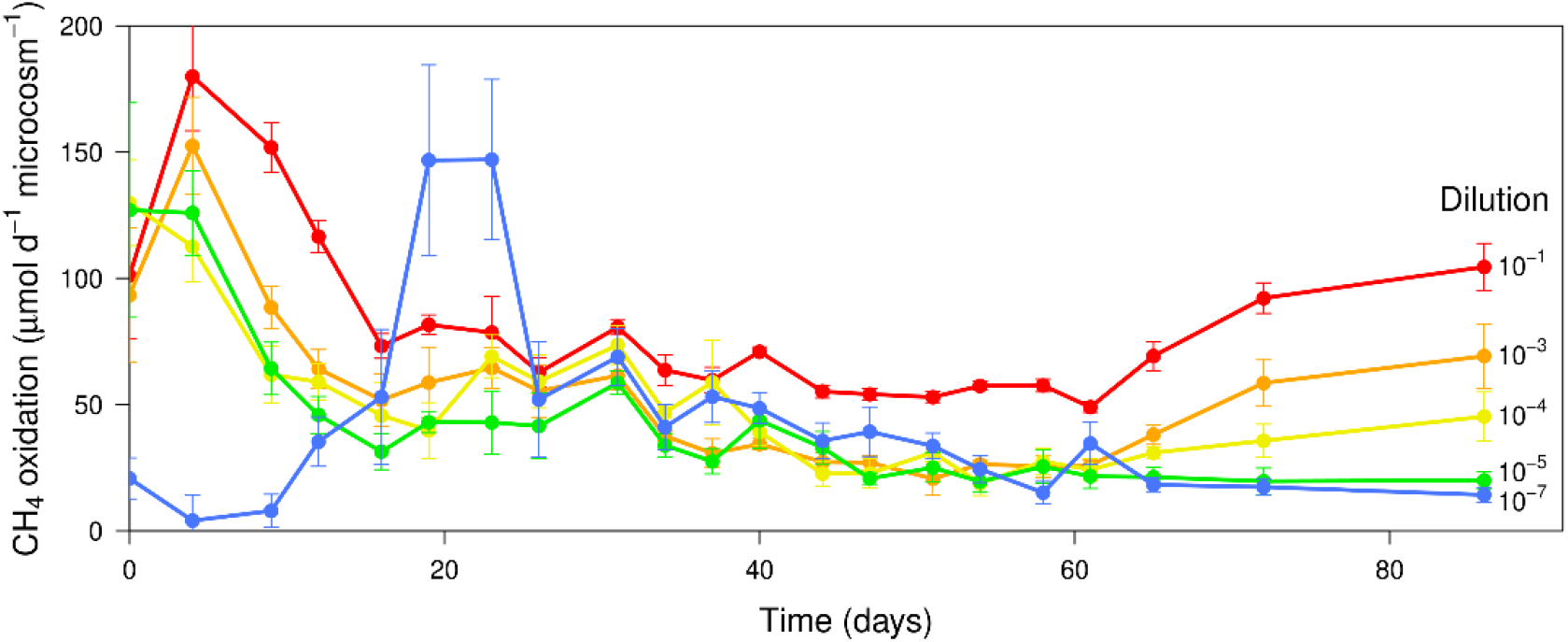
Net headspace CH_4_ consumption rates of microcosms as a function of dilution level and sampling date. Data of the two temperature treatments were combined because no temperature effect was detected. Symbols show mean ± se (n=6).

## Discussion

Using a dilution-to-extinction approach, we demonstrate that diversity loss can affect net methane (CH_4_) consumption by methanotrophic communities extracted from a landfill cover soil. Methanotrophic bacteria perform critical biogeochemical functions in many ecosystems; in landfills, including the one we studied here, methanotrophs typically capture nearly all CH_4_ produced during the decomposition of waste material [34]. Landfills are an important anthropogenic source of CH_4_. Establishing and managing cover soils so that they efficiently filter CH_4_ is a key tool to mitigate landfill CH_4_ emission. Our experiments suggest that this methanotroph-driven ecosystem service critically depends on the presence of a sufficiently diverse methanotrophic community.

Experimental research has shown that typical BEF relationships in communities of higher organisms such as plants generally are positively decelerating, i.e. the benefit of additional species decreases with the diversity of the system. These relationships may even saturate because of functional redundancy since the loss of a few species will functionally be compensated by others with no resulting net effect on community-level function [37]. It has been argued that microbial communities are likely to exhibit pronounced functional redundancy [38] due to their tremendous diversity and due to the relative simplicity of microbes compared to plants [21–23]. If this was true, microbial diversity loss would only lead to a loss of function at the lower end of the diversity gradient [39]. These considerations are highly relevant for methanotrophic communities, for several reasons. First, methanotrophs all perform the same specialized metabolic function that is restricted to comparably narrow phylogenetic groups. Second, methanotroph diversity typically is relatively low with rarely more than a few dozen OTUs. Third, methanotroph diversity and activity are vulnerable to disturbance, for example by high mineral N levels as they can result from fertilizer application [40]. A total of 58 methanotrophic OTUs were detected in the landfill cover soil under investigation. Although this number might appear high, we observed a clear reduction of CH_4_ consumption rates under dilution-induced diversity loss, contrasting the notion that functional redundancy protects ecosystem functioning from diversity loss. Importantly, these detrimental effects manifested despite the fact that the OTUs eliminated by dilution were of low abundance. This finding corroborates earlier studies that used isotope probing of methanotrophic communities and found that subordinate taxa contributed significantly to community-level functioning [41].

Serial dilutions that are used to inoculate sterile soil microcosms are the method of choice to reduce diversity in microbial BEF studies where the number of species is high and their identity unknown [42, 43], and allows to the preferential extinction of rare taxa [35]. In our study, the applied serial dilutions were highly correlated with OTU-based diversity metrics for methanotrophic and non-methanotrophic bacterial communities, but the 10^−1^ dilution deviated from the overall pattern. In this dilution (but not in others), a milky residue and gel-like pellets occurred during DNA extraction and likely reduced DNA yields and led to a relative underestimation of diversity. The primary treatment applied, dilution level, was unaffected by this artifact and explained the observed responses better that the OTU-based metrics unless the least-diluted treatment was excluded. Irrespective of this methodological caveat, the relationship between diversity and methanotrophic activity was robust, resembling patterns of BEF-relationship found in plant communities [44].

In contrast to diversity experiments in which artificial communities are assembled *de novo* from isolated strains [45], dilution treatments decrease the abundance of both target and non-target microorganisms, and our study was no exception. However, soil communities recovered well during the 4-week preincubation, with no remaining differences in total bacterial and only relatively small differences in methanotrophic community size at day zero except for the most diluted community which required more time to recover. Non-methanotrophic bacterial community size decreased over the course of the experiment, but this change was independent of dilution. Methanotroph community size showed a dilution-dependent temporal dynamics, with more diluted samples displaying higher initial growth rates. This likely was the case because methanotrophic abundances were below the carrying capacity of the system, possibly combined with reduced competition for oxygen and nutrients by heterotrophs. As a result, methanotrophs initially grew exponentially and community size overshot in microcosms of the highest dilution treatment. A similar effect has been observed in other studies, including sewage microbial communities [46] and grassland denitrifier communities [47], in which the highest dilutions showed the strongest initial growth in bacterial abundance. However, in our study, this effect was transient, with trends in community size that reversed towards the end of the study. Despite the temporal dynamics in community size, effects of dilution and OTU diversity on CH_4_ consumption remained remarkably stable, indicating that the BEF-effects found were unrelated to the initial fluctuations in community size.

Towards the end of the experiment, net CH_4_ consumption increased and biodiversity effects became most pronounced. Many studies with plant systems have shown that BEF relationships grow larger through time [48–52]. The methanotrophs in our study thrived on above-atmospheric CH_4_ concentrations, displayed an apparent low-affinity kinetic [53], and had a relatively short generation time compared to plants; our three-month experiment under near-optimal growth conditions will thus have allowed for many generations. BEF-effects that grew through time also have been observed in microbial systems [54, 55], although patterns generally varied among environments. We did not investigate the specific mechanisms that promoted biodiversity effects; however, there is evidence that stronger BEF relationship emerge from the evolution of less negative species interactions [54] or from an increase in interspecific niche complementarity (microbes: [56]; plants: [57]).

Methanotroph diversity may buffer ecosystem functioning during environmental fluctuations [58–60]. Methanotrophs isolated from environmental samples typically exhibit distinct temperature optima. We therefore expected stronger BEF relationships in the model communities that were exposed to a diurnal temperature cycle because we reasoned that more diverse communities would span a wider range of temperature optima. However, we did not find such an effect. One reason may be that we assessed CH_4_ consumption outside the incubator, under constant temperature, regardless of the original treatment. We nevertheless would have expected that effects present in the incubator would have persisted for some time and therefore would have been detectable in the measurements. Interestingly, we also did not observe any effect of temperature regime on the diversity of methanotrophs, their specific community composition, or their growth. This implies that these communities remained unaffected by temperature cycling, possibly because the temperature variability resembled the fluctuations found naturally in the original landfill soil and species or entire communities were adapted to these conditions.

The food web in our microcosm communities was largely CH_4_-based, with no plants present in the microcosms and plant cover was relatively low at the original landfill sampling site. However, our dilution-to-extinction approach not only reduced the diversity of methanotrophs but also the one of non-methanotrophic bacteria (and potentially also of other groups of organisms that we did not measure, for example protists). In natural systems, the diversity of even remote taxa often co-varies [10, 61], so that BEF relationships observed in such systems reflect the compound effects of diversity changes across multiple, potentially interacting, taxonomic groups. Methanotrophs require other microorganisms for sustained growth [60, 62], most likely to remove metabolites that otherwise accumulate to inhibitory levels. Recent experiments also have shown that volatiles produced by heterotrophs, specifically dimethyl sulfides, can promote methanotrophic growth even if heterotrophs are physically isolated [63]. While we think that the reduced methanotrophic activity resulted primarily from a loss of methanotrophic strains, these findings raise the possibility that a loss of interactions with heterotrophic bacteria further exacerbated the reduced functioning in low-diversity communities.

Methanotrophs play a key role in the regulation of climate by oxidizing large fractions of the CH_4_ produced in anaerobic environments, thereby preventing CH_4_ emissions. Methanotrophs also oxidize significant amounts of atmospheric CH_4_ [2]. Our results indicate that a reduction in the diversity of methanotrophs can affect system-level CH_4_ consumption, through mechanisms independent of community size. The past years have seen an accelerated rise in atmospheric CH_4_ loads [3], which underlines the critical importance of diverse methanotrophic communities in sustaining stable CH_4_ oxidation services in managed and natural wetlands. More generally, our study suggests that microbial diversity supports microbially-driven ecosystem functions in a way similar to the one found in plants, i.e. positive BEF-relationships appear universal across very different taxa and spatial scales.

## Acknowledgements

This project was funded by the Swiss National Science Foundation (grant 144065 to PAN) and the University of Zurich. We thank René Husi, Thomas Rime, and Mariela Soto Araya for help with laboratory analysis. We acknowledge the contribution of the staff at the McGill University and Génome Québec Innovation Center, Montreal, Canada, for the sequencing service. We further are indebted to Kentaro Shimizu and the Genetic Diversity Centre (GDC), ETH Zurich for generous help with laboratory facilities. PAN acknowledges support from the University of Zurich Research Priority Programme Global Change and Biodiversity.

## Methods

### Soil sampling

We collected soil from a landfill (Liestal, Basel-Land, Switzerland, 570 m a.s.l. 47°29′N, 7°45’E, [64]) that had accumulated c. 3·10^6^ m^3^ of household, office and construction waste before it was covered with 2 m of soil in the 1990s. The cover soil is a heterogeneous mixture, loamy in texture (40–60% silt), slightly alkaline (pH 7.3–7.9), and contains pebbles, rocks, boulders and construction material. The annual temperature at the site averages around 9°C and annual precipitation around 1000 mm. Active methanotroph communities are dominated by type Ia methanotrophs [65]. We collected soil from a depth of 30— 50 cm (ca. 150 cm west of sampling location C1 described in [33]), sieved the samples (2 mm mesh) and stored them at 4°C. Before the start of the experiment, the soil was incubated for two weeks at room temperature under an atmosphere of air spiked with 1% CH_4_ (v/v) to re-activate the methanotrophs.

### Experimental setup

Inocula were prepared by homogenizing 10 g of the collected landfill soil in 20 mL of sterile H_2_O with a pestle and a mortar. This slurry was then repeatedly diluted 10—fold by mixing 1 mL of the highest dilution with 9 mL of sterile H_2_O until a final dilution of 10^−10^ was obtained. This procedure was repeated three times with three separate original soil subsamples, resulting in three independent dilution series. From each dilution, 9 mL were transferred to an Erlenmeyer flask with 1 mL NMS [66] and 40 g γ-sterilized soil (collected at Nenzlingen, Switzerland; 47°33’N, 7°34’E, 520 m a.s.l.; silty clay loam soil, 41% clay, 52% silt 3.9% C, 0.33% N, pH ≈ 7,8; [67]). This matrix soil has very low natural methanotroph abundances, which minimizes interference from remaining soil DNA (the relative abundance of the *pmoA* gene measured by qPCR in this soil was less than 0.01% of that measured later in inoculated microcosms). The flasks were closed with gas-permeable cotton stoppers and incubated in air-tight jars for one month to allow methanotrophic communities to recover. Headspace CH_4_ concentrations were brought to 1% by injecting CH_4_ with a syringe. The flasks were ventilated regularly. Headspace CH_4_ was monitored by gas chromatography and additional CH_4_ was injected when needed. For the experiment, we used a total of five dilutions: four that showed a recovery of methanotrophic activity during the pre-incubation (dilutions 10^−1^, 10^−3^, 10^−4^, 10^−5^) plus one that did not show any significant methanotrophic activity at the beginning of the study (dilution 10^−7^) but that recovered later. The experimental microcosms were created by transferring 2 g of cultured soil with an additional 2 g of sterile soil matrix to 50 mL centrifugation tubes (Sarstedt, Nümbrecht, Germany).

### Experimental conditions

We incubated each of the 15 distinct communities together with negative controls (sterile soil only) in separate incubators with temperature averages of 15°C (3 dilution series × 5 dilutions × 7 replicates + 2 controls = 107 microcosms per incubator). In the first incubator, the temperature was kept constant. In the second, the temperature ramped up from 10 to 20°C in 12h, and then down again to 10°C in another 12h. Within the incubators, the microcosms were kept in air-tight chambers that were connected to a custom-built system that measured and controlled CH_4_ concentrations. Headspace air was pumped through a heat exchanger that equilibrated the gas to room temperature. Then, the sampled air was dried in a tube filled with silica gel beads before CH_4_ concentrations were determined in a CH_4_ detector cell (TGS 2611, Figaro Inc., Arlington Heights, IL). Whenever CH_4_ concentrations fell below approximately 6000 ppm, a mixture of 5% CH_4_ and 10% O_2_ in N_2_ was added through a solenoid valve. The readings obtained from the semiconductor gas sensor were not of very high precision. We therefore used the same sensor and electronics to control CH_4_ concentrations in both incubators (switched between incubators with a solenoid valve). The concentration therefore did not differ systematically between the treatments. CH_4_ concentrations were further monitored by taking headspace samples from the boxes with a syringe 1–2 times a day. CH_4_ concentrations averaged 7400 μmol CH_4_ mol^−1^ throughout the experiment (gas chromatographic analysis).

### CH_4_ consumption and CO_2_ release rates

To determine net CH_4_ and CO_2_ exchange rates, replicate microcosms of all 30 communities were removed from the incubators and placed in air-tight 3 L jars with elevated CH_4_ concentrations. Headspace samples were collected with a syringe and analyzed for CH_4_ and CO_2_ by gas chromatography (Agilent 7890N gas chromatograph; CH_4_ was detected with a flame ionization detector; 12’ Porapak Q column; isothermic at 80°C; He carrier gas; CO_2_ was determined on the same detector after reducing CO_2_ with H_2_ on a Ni-catalyst; Agilent Technologies Inc., Santa Clara, CA). A total of 3 headspace samples were collected over 3 days, except when CH_4_ consumption rates were low, in which case 3–4 samples were taken over 4 days. Then, microcosms were placed back in the incubators. Gas exchange rates were calculated by linear regression of headspace concentrations against sampling time, with consumption rates converted to μmol microcosm^−1^ d^−1^ by applying the ideal gas law.

### DNA extraction

After 0, 31, 58, and 86 days of incubation, one of the replicates of each of the 15 dilution × dilution series combinations was removed from both incubators and DNA retrieved using the xanthogenate-based extraction described in [45]. Extracted DNA was purified twice using the PowerClean Pro DNA Clean-Up Kit (MoBio, Carlsbad, CA) to remove PCR inhibitors.

### Quantitative PCR

We quantified the size of total bacterial communities and methanotrophic communities by determining the abundances of the 16S rRNA gene (primer 27F and 1406R) and of the methanotrophic *pmoA* gene (subunit A of the particulate methane mono-oxygenase gene; primers A189F and mb661) by quantitative PCR (StepOne real-time PCR system, Applied Biosystems, Foster City, CA; SI Tables S1, S2). For calibration, a serial dilution of purified DNA from *Methylococcus capsulatus* Bath (quantified with a Qubit Fluorometer, Invitrogen, Carlsbad, CA) was included in duplicate in each run. Additionally, we included reference samples with DNA of *M. capsulatus* on all plates and then standardized between plates using the geometric mean [68].

### Sequencing

The composition of total bacterial and methanotrophic communities was determined by amplifying in duplicate the variable regions V1 and V2 of the 16S rRNA gene and the *pmoA* gene (primer pairs 27F/341R and A189F/mb661, respectively; SI Tables S1 and S2). The duplicate PCR products were pooled and purified with the GeneJET PCR purification kit (Thermo Scientific, Waltham, MA) and quantified fluorometrically using the Qubit dsDNA BR Assay Kit (ThermoFischer, Waltham, MA) on the Spark 10M Multimode Microplate Reader (Tecan, Männedorf ZH, Switzerland) with a standard curve (0–100 ng DNA μL^−1^). Samples were barcoded with the Fluidigm Access Array technology and the amplified regions were paired-end sequenced on the Illumina MiSeq v3 platform at the Genome Quebec Innovation Center, Montreal, Canada. Forward and reverse sequence data were merged, quality controlled and clustered by OTU using a customized pipeline [45] that based on the algorithms implemented in USEARCH v9 [69]. All *pmoA* OTU sequences were blasted against the NCBI database and sequences that did not yield a match or were not *pmoA* were removed. For taxonomic classification, the OTU centroid sequences were mapped using the naïve Bayesian classifier implemented in MOTHUR [70] with a minimum bootstrap support of 60%. 16S_V1-V2_ and *pmoA* sequences were mapped against sequences from the SILVA database [71], version 123, retrieved from https://www.arb-silva.de, and the *pmoA* taxonomic classifier [72], respectively.

### Alpha diversity and phylogenetic diversity

Phylogenetic trees (UPGMA) for the 16S rRNA gene and the *pmoA* gene were calculated with Clustal Omega [73] using the web services from the EMBL-EBI [74]. *pmoA* phylogenies were built by adding sequences retrieved from GeneBank for all accessions in the *pmoA* phylogeny in Fig. 1 in [75]. The phylogenetic diversity (PD) of each sample was calculated as the total branch length of the phylogenetic subtree defined by the strains found in each sample (function calcPD; https://github.com/pascal-niklaus/pdiv). We randomly subsampled both 16S and *pmoA*-based OTU tables 5000 times to an even depth (10000 sequences 16S rRNA, 18000 sequences *pmoA)* and determined OTU richness (S), the Shannon index of OTU distribution (H), and phylogenetic diversity (PD) of these rarefied OTU sets. Rarefaction reduced S on average by 7% and 25% for *pmoA* and 16S, respectively. H was nearly invariant to rarefaction, with reductions < 1%. Reductions in PD averaged 4% and 22% for *pmoA* and 16S, respectively. A few samples had too few sequences for rarefaction; this was the case in particular for *pmoA* in the 10^−7^ dilution at day 0. These samples are marked in figures and were excluded from statistical analyses.

### Statistical analysis

We used analysis of variance (ANOVA) based on general linear models (R 3.5; http://r-project.org) and mixed models (ASReml 3.0; VSN International, Hemel Hempstead, UK) to test for effects of our experimental treatments on methanotrophic activity and community size. Fixed effects were the experimental treatments dilution (*log*-transformed), temperature (two-level factor), and their interactions. Community composition (the 15 specific combinations of dilution series and dilution) was included as random effect. Since the goal of the dilution treatment was to reduce the diversity of methanotrophs, we fitted additional models with OTU diversity as explanatory variables. For biodiversity effects, the null hypothesis is that contributions of OTUs to functioning are additive, i.e. that OTUs do not interact in a systematically positive (or, as rarely found, negative [76]) way. Testing for diversity effects by regression thus requires that data remain untransformed. This precludes any data transformation to address heteroscedastic residuals because this could introduce spurious false positives (see [77] for details). In our analyses, residuals generally were normally distributed and homogenous except for the 10^−7^ dilution which sometimes showed enhanced variation; we accounted for this heteroscedasticity by fitting a heterogeneous random term (‘idh’ option of ASReml with a separate, larger, variance component for dilution 10^−7^). For the analysis of qPCR data, we adjusted for heteroscedasticity by weighing data with the inverse of the within-dilution level variance. Effects of temperature and dilution treatments on community composition were tested using permutational ANOVA [78] as implemented in the ‘adonis’ function of R-library ‘vegan’. To test for effects of the temperature treatment, data were permuted among temperature treatments within combinations of dilution level and series. To tests for effects of dilution, dilution levels were permuted within dilution series, keeping pairs of microcosms with different temperature treatment as unaltered unit of randomization. These tests were performed at different levels of taxonomic resolutions, from OTUs to genera to families to classes to subphyla to phyla, and test results adjusted to a false discovery rate of 5% using the method of Benjamini and Hochberg [79] as implemented in the R function ‘p.adjust’. We determined Bray-Curtis dissimilarities among communities based on square root-transformed abundance data and used principal coordinate analysis (PCoA) to map these differences (β diversity) into two-dimensional space using the R function ‘cmdscale’. These ordination plots were used to inspect the trajectories that dilution and time-series delineated in composition space, and whether these trajectories depended on the applied temperature treatment.

