## Supplementary Material for "Experimental erosion of microbial diversity decreases soil CH_4_ consumption rates"

### Supplementary Information

**Table S1 |** List of reagents and their concentration in the solution mix as well as the primer sequences used to amplify 16S rRNA and *pmoA* genes for qPCR and for Illumina sequencing (including the Fluidigm adapter sequences CS1 and CS2, in bold).

**PCR for sequencing (50 µL volume)**

| | |
|---|---|
| 25 µL | NEBNext Q5 Hot Start HiFi PCR Master Mix (New England Biolabs, Ipswich, MA) |
| 1 µM | Forward primer |
| 1 µM | Reverse primer |
| 15 µL | DNA |
| 16S rRNA gene: | |
| Forward primer<br>(Lane, 1991) | CS1-27F (5'– **ACA CTG ACG ACA TGG TTC TAC A**AG AGT TTG ATC CTG GCT CAG –3') |
| Reverse primer<br>Reverse complement of 341f (Muyzer *et al.*, 1993) | CS2-341R (5'– **TAC GGT AGC AGA GAC TTG GTC T**CT GCT GCC TCC CGT AGG –3') |
| *pmoA* gene: | |
| Forward primer<br>(Holmes *et al.*, 1995) | CS1-A189F (5'– **ACA CTG ACG ACA TGG TTC TAC A**GG NGA CTG GGA CTT CTG G –3') |
| Reverse primer<br>(Costello and Lidstrom, 1999) | CS2- mb661(5'– **TAC GGT AGC AGA GAC TTG GTC T**CC GGM GCA ACG TCY TTA CC –3') |

**qPCR (20 µL volume)**

| | |
|---|---|
| 10 µL | KAPA SYBR FAST qPCR Master Mix (2x) ABI Prism (Kapa Biosystems, Wilmington, MA) |
| 0.2 µM | Forward primer |
| 0.2 µM | Reverse primer |
| 8.2 µL | $H_2O$ |
| 1 µL | DNA |
| *pmoA* gene: | |
| Forward primer<br>(Holmes *et al.*, 1995) | A189F (5'– GGNGACTGGGACTTCTGG –3') |
| Reverse primer<br>(Costello and Lidstrom, 1999) | mb661(5'– CCGGMGCAACGTCYTTAC –3') |
| 16S rRNA gene: | |
| Forward primer<br>(Lane, 1991) | 27F (5'– AGAGTTTGATCCTGGCTCAG –3') |
| Reverse primer<br>(Stahl *et al.*, 1988) | 1406R (5'– ACGGGCGGTGWGTRC –3') |

**Table S2 |** PCR cycling conditions for amplifying 16S rRNA and *pmoA* genes for sequencing and qPCR.

| **PCR** | **Time (s)** | **Temp. (°C)** |
|---|---|---|
| Initial denaturation | 30 | 98 |
| 20 cycles (16S rRNA) or | | |
| 25 cycles (*pmoA*) of: | | |
| Denaturation | 10 | 98 |
| Annealing | 30 | 55 |
| Extension | 30 | 65 |
| Final extension | 300 | 65 |

| **qPCR 16S rRNA** | **Time (s)** | **Temp. (°C)** |
|---|---|---|
| Initial denaturation | 180 | 95 |
| 40 cycles of | | |
| Denaturation | 15 | 95 |
| Annealing | 30 | 55 |
| Extension | 30 | 72 |
| Data collection | 30 | 85 |

| **qPCR *pmoA*** | **Time (s)** | **Temp. (°C)** |
|---|---|---|
| Initial denaturation | 180 | 95 |
| 7 cycles of | | |
| Denaturation | 15 | 95 |
| Annealing | 30 | 62 (-1/cycle) |
| Extension | 30 | 72 |
| 33 cycles of | | |
| Denaturation | 15 | 95 |
| Annealing | 30 | 55 |
| Extension | 30 | 72 |
| Data collection | 30 | 85 |

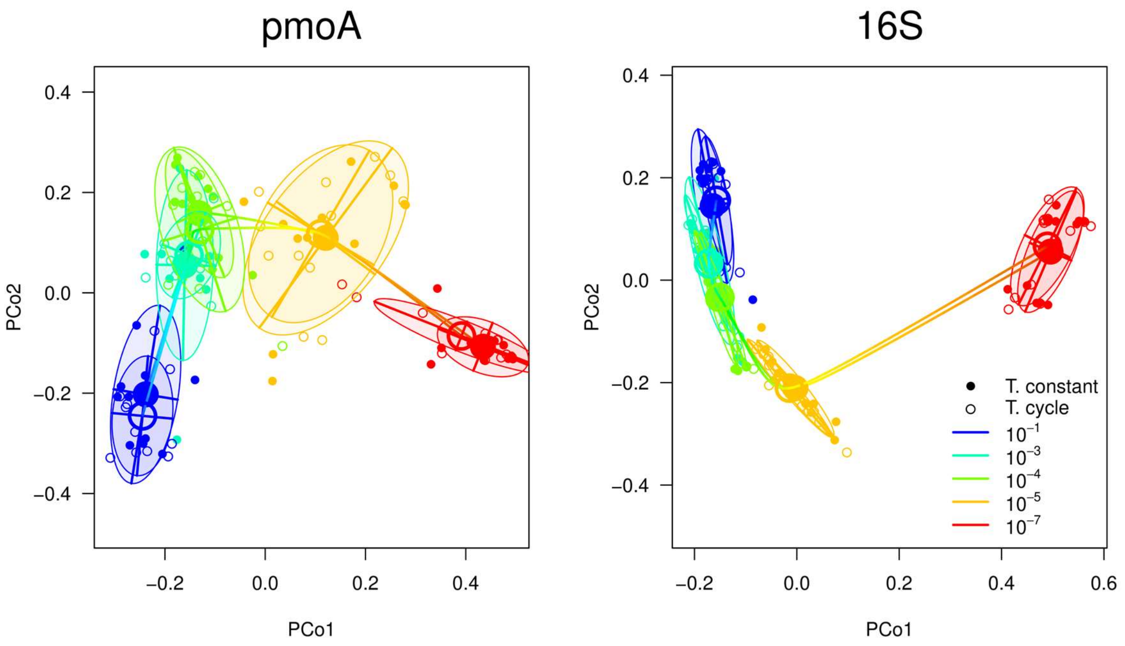


**Figure S1 | Community dissimilarities among the different dilution and temperature treatments.** Ordination plots show principle coordinates of Bray-Curtis dissimilarities calculated from square-root transformed *pmoA* and 16S sequence abundances. Ellipsoids indicate 67% confidence intervals.
